# Increased ‘selfness’ in the tumor emerges as a possible immune sculpting mechanism: A pan-cancer data analysis of 32 solid tumors in TCGA

**DOI:** 10.1101/2024.03.18.585489

**Authors:** Naren Chandran Sakthivel, Anoushka Chinmayi, Nagasuma Chandra

## Abstract

Tumors pose a unique challenge to the immune system since they straddle the boundary between ‘self’ and ‘non-self’. T-cells recognize tumors that contain ‘non-self’ neo-antigens. They can also recognize tumors that contain aberrantly expressed self-antigens, highlighting the importance of the central tolerance and the tuning of the T-cell repertoire in the thymus. Therefore, the similarity to the thymic expression profiles must have information in it to influence the T-cell repertoire and what self-peptides are recognized. We utilize this principle in a pan-cancer analysis and develop a thymus-like or ‘selfness’ score (TLS) based on the gene-expression similarity to thymi, indicative of recognizability of tumors by T-cells. We show that the TLS is indicative of patient survival in 8 different TCGA cohorts, indicating gene expression modulation to mimic that in thymi as a potential immune sculpting mechanism. Surprisingly, we also see an inverse relationship between TLS and the degree of immune infiltration.

## Introduction

The components of the immune system exhibit an intricate interplay with various cancers, shaping the evolutionary trajectory of tumors throughout tumor formation and progression and defining disease outcome. The process is broadly described as immunoediting and consists of three sequential phases - elimination, equilibrium, and escape and sits at the interface between initial immune responses, ongoing immune surveillance, and the phenomenon of immune sculpting that lead to the development of immune evasion mechanisms by cancer cells^1^. Known mechanisms of immune evasion by tumors include altering antigen expression, modifying antigen recognition by downregulation of class-1 HLA molecules to reduce recognition of tumor antigens^2^, creating immune-privileged sites by creating an immunosuppressive environment by attracting a larger number of Tregs or secretion of immunosuppressive cytokines or upregulation of immune checkpoint molecules^3^, immune exclusion by limiting immune cell infiltration into the tumor microenvironment^4^ or by induction of immune anergy^5^, mimicking self-tolerance signals and exploiting tolerance mechanisms, tumors can navigate the immune landscape without triggering an aggressive response. The self versus non-self discrimination is a pivotal point in defining immune recognition and tumor cell killing. Subverting the distinction between self and non-self would provide an effective mode of immune evasion to the tumor cells. Not much is known about what such mechanisms are or whether such mechanisms even exist.

The self/non-self paradigm itself has been well studied and is known to be established over all T-cells by a stringent filtering process in the thymus. Immature T-cells entering the thymus first undergo positive selection in the cortical region: T-cells that cannot bind to pMHC complexes undergo apoptosis. The surviving contingent of T-cells then moves onto the medullary region and experiences negative selection, wherein T-cells that strongly bind to pMHC complexes with self-antigens are eliminated. The surviving T-cells comprise the mature but naive T-cells available to elicit immune responses^6,7^. Therefore, it is the negative selection that ensures the removal of potentially autoreactive T-cells. If T cells are to be trained against recognition of self-antigens, then by necessity, the self-antigens must be present in the thymus for the training process. Medullary thymic epithelial cells (mTEC) are the primary source of self-antigens for the negative selection process, expressing tissue-restricted antigens (TRA) from throughout the body. The lack of a protein’s expression in these cells is causally linked to the generation of self-reactive T cells targeting peptide antigens derived from that protein^8^. Expression of TRA-encoding genes is limited to a mature subset of mTEC^hi^ cells^9^. AIRE and FEZF2 are two transcription factors known to regulate this process^10,11^. Available data suggests that each individual mTEC^hi^ cell expresses only a small fraction of the proteome, but the aggregate of all such cells covers nearly the entire proteome^12^, making the training of the ‘self’ comprehensive.

A logical next question is how a tumor gets perceived as non-self and how it can evade such recognition? T-cells can recognize specific peptide antigens from the tumors that arise due to mutations, indels, and structural variations, forming neo-antigenic peptides, that are presented by the Human Leukocyte antigens (HLA) molecules^13,14^. Only a small fraction of the proteome is presented on HLA molecules at any given time, and the odds of any specific genetic alteration resulting in the production of neo-antigens is therefore limited. In addition to neo-antigens, other tumor specific antigens include genes selectively expressed in tumors and absent in most normal tissues^15–17^. These specific antigens are termed as tumor specific antigens and T-cell response against them are expected to be specific only to tumors since the other tissues do not express or present these antigens. However, there are several instances of T-cells reacting against tumors using what are ostensibly self-antigens, termed as Tumor Associated Antigens (TAA)^18–20^.

Since the recognition of a tumor cell as ‘non-self’ by a T lymphocyte is determined by the gene expression pattern in mTEC^hi^ cells during the developmental phase, we hypothesized that a possible option for a tumor is to evade immune responses is by sculpting its gene expression pattern. We systematically investigate this in the current study and test T-cell recognizability of tumors in a large-scale analysis of TCGA cohorts, by utilizing transcriptomic data. To enable this, we develop a ‘Thymus-likeness score (TLS)’, which is based on the similarity of global gene expression profiles of tumor cells with that of mTEChi cells. We show that stratification of tumors by this score is indicative of tumor microenvironment composition and survival differences across several cancer types. Put together, we observe that in general, tumors with lower TLS i.e. the ones that are geared for better recognizability are the ones that are least infiltrated. The paradoxical inverse relationship between recognizability and immune infiltration suggests the presence of different routes to immune evasion of tumors - with highly recognizable tumors evading destruction due to insufficient immune machinery, and the rest enacting evasion due to being poorly recognizable. Our results imply the presence of an immune sculpting mechanism for increasing ‘selfness’ of a tumor and a compensatory mechanism that drives tumor infiltration, together influencing the disease outcome.

## Results

The primary goal of our analysis was to establish a quantitative view of the recognizability of tumors from a T-cell perspective, and explore prognostic and molecular insights that arise from inspecting tumors under this lens. Specifically we investigated if tumor transcriptomes are tuned to become less ‘non-self’, so as to evade recognition by CD8 T cells, and the associated consequences of such a phenomenon. To assess this, we develop a thymus-likeness score to model the tumor recognizability and design a robust pipeline to examine the links between this score and immune infiltration and cancer prognosis.

### Formulating the ‘ Thymus-likeness score’ (TLS)

We first develop a quantitative measure of ‘selfness’ to estimate the recognizability of tumors and characterize it across 10810 samples from 32 solid tumor cohorts in The Cancer Genome Atlas and 54 normal tissues in the Human Protein Atlas. We define ‘selfness’ from the perspective of central tolerance based on thymus likeness, which is the extent of similarity in the overall expression profile of a sample to a representative value of mTEC^hi^ cells and compute the Spearman Correlation coefficient to output a score that we refer to as the ‘Thymus-likeness score’ (TLS). Being a correlation coefficient, the TLS conveniently maps between −1 and 1 (both inclusive), is easy to interpret, has good computation speed, and is robust to outliers. As a corollary, the ‘non-selfness’ can be computed as the sign inversion of the selfness score (−1 * TLS). A highly positive TLS indicates high similarity in the relative ordering of the gene expression values of the query sample with respect to the mTEChi cells. To derive the reference profile for the mTEChi cells, we use gene expression profiles from two publicly available datasets^21,22^. These two datasets comprise eight mTEChi samples isolated from pediatric thymi. Partial or complete thymectomy is common in pediatric cases to facilitate the operation on the heart^23^, which is the category of samples used to isolate these cells in the chosen datasets. No such datasets were available for samples isolated from adult thymi. Since the thymic output is maximum after birth and the naive T-cell population in adults is primarily maintained by replication of previously existing peripheral naive T-cells^24,25^, pediatric thymi constitute an appropriate tissue source.

To validate the scoring scheme, we tested if the TLS captures known similarities of different tissue types with the thymus. We computed the TLS for 43 normal tissues, for which data was available publicly from the Human Protein Atlas^26^ consensus RNAseq dataset (Figure 1A). We find the scores to be in the 0.6 to 0.8 range (median ∼0.78), and reassuringly the thymus itself exhibiting the highest TLS, followed by the tonsils, which are known sites of extrathymic T-cell maturation^27^ (Fig 1A). Testes, brain, retina and spinal cord had the lowest TLS scores, consistent with a previous report, which showed that mTEC^hi^ cells in mice express fewer tissue-specific genes from such immunoprivileged sites^28^. As a secondary check, we also compared the tissue expression profiles among themselves in an all-vs-all manner, which showed high similarity among similar tissues and clustering of the aforementioned low TLS tissues, further demonstrating the biological significance of the use of gene expression profiles for comparing different tissue types (Supplementary figure S1C).

**Figure 1.**
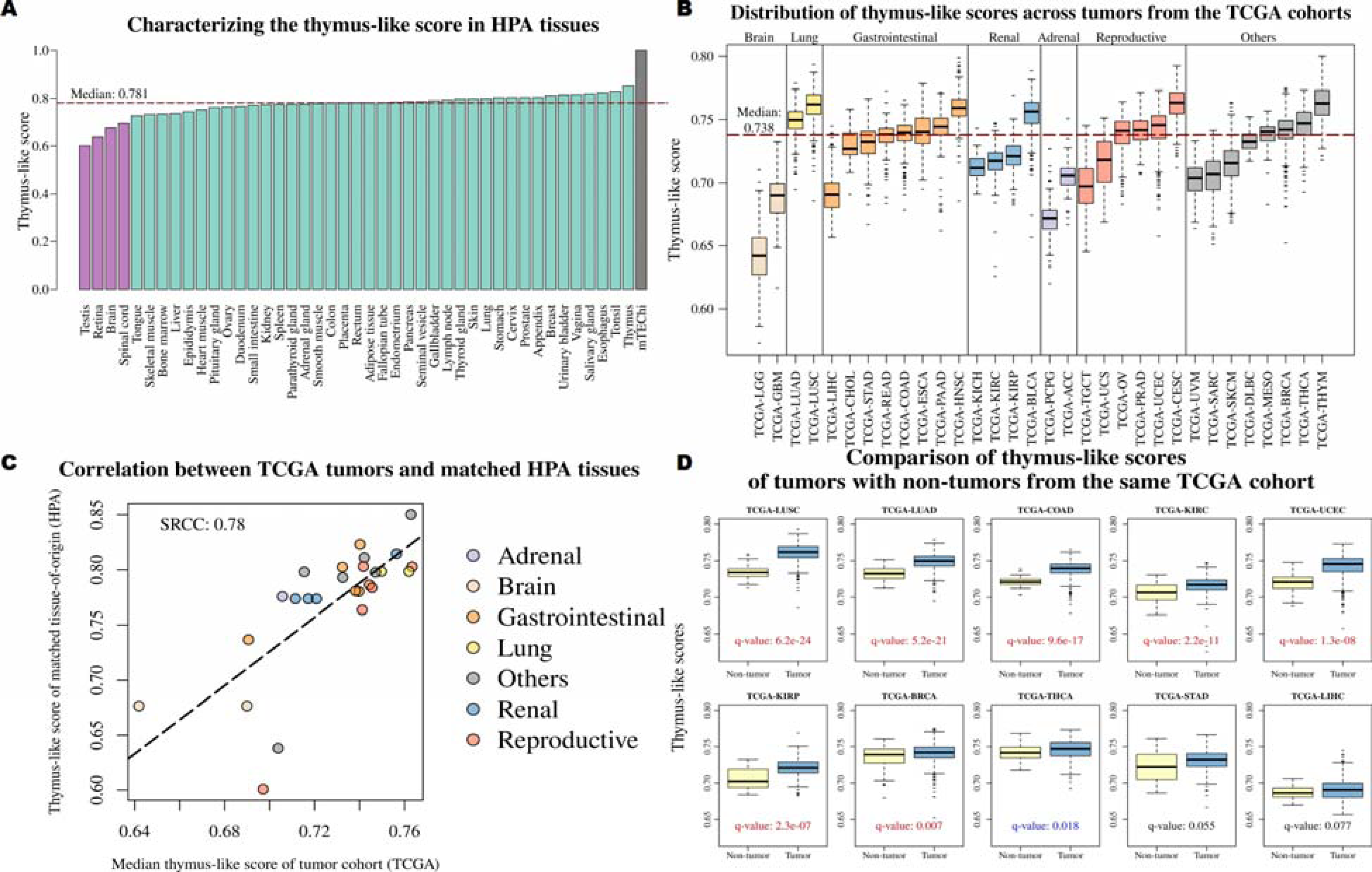
Characterizing the “thymus-like” score across physiologically normal and tumor tissues. **(A)** The thymus-like score of tissues from the Human Protein Atlas. Univariate segmentation analysis results in two broad clusters, marked with different colors here. **(B)** The distribution of thymus-like scores across the 32 solid tumor datasets from TCGA. The datasets have been arranged by organ/organ system of origin. The dotted line indicates the median TLS across all samples. **(C)** Comparison of median TLS of different TCGA cohorts with scores of matched normal tissues from the Human Protein Atlas. **(D)** Comparison of thymus-like scores of tumors with adjacent non-tumor samples from TCGA cohorts with at least 20 non-tumor samples. Significance values displayed in each subplot were calculated using Wilcoxon rank sum test and then adjusted for multiple hypothesis correction using False Discovery Rate. The ten cohorts where the tumors show higher TLS are showcased here

### Characterizing the TLS for tumors from The Cancer Genome Atlas

Next, we calculated the TLS for 10810 samples from 32 solid tumors from The Cancer Genome Atlas (TCGA) project^29^. The TLS of a sample is in a scale of −1 (non-self) to 1 (self) and the lower it is, the more its non-selfness and as a corollary, higher the probability that it can be recognized by T cells. Overall the TLS ranged between 0.6 to 0.8 (Figure 1B), with considerable variation across tumor types. Like with the HPA tissues, the lowest value of TLS is still ∼0.6, indicating that tumors with the lowest TLS still have good concordance between the expression profile of the mTEC^hi^ cells. Next, we compared how adjacent non-tumor samples fared in 13 cohorts with sufficient number of non-tumor samples (n ≥20). Tumors showed significantly increased TLS (Wilcoxon rank sum q-value < 0.05) in eight of the thirteen datasets (Figure 1D). These non-tumor samples are tumor-adjacent and may represent an intermediate state between the physiologically normal and cancer conditions. Our results indicate that tumors attempt to become more ‘self’ or thymus-like to evade recognition by the adaptive immune system. This observation is consistent with the immunoediting theory that the attack of tumors by immune cells results in the selection of immune-evasive tumors.

Interestingly, just as with HPA, we found that tumors that arise from thymic tissue (THYM cohort) have the highest TLS score, while those tumors that originate from the brain (LGG and GBM cohorts) and testes (TGCT cohort) have low TLS values. We reasoned therefore that the score could be dependent in part on the tissue of origin, as different tumors have different prognosis based on the immune responses the tissue type can elicit. To test this hypothesis, we compared median TLS from TCGA datasets against scores calculated for physiologically normal tissues in the Human Protein Atlas that best matched the tumor’s tissue of origin and found that they indeed correlated significantly (Figure 1C). This indicates that a tumor’s TLS is potentially set by its tissue of origin, and that the context of the tumor’s origin is crucial in determining how easily recognizable the tumor will turn out to be. Within each tumor type, however, we observe a wide variation of the TLS score for different patient samples, which indicate that they may elicit different extents of immune response and disease outcomes. At this juncture, we have no information about the nature of the mathematical relationship between the TLS and these outcomes, so we devise a robust pipeline to examine the same, which we discuss further in the next section.

Apart from these aforementioned characteristics, we also noticed an interesting trend regarding TLS and the rarity of the cancer. Of the 32 TCGA solid tumor cohorts analyzed, ten cohorts are for rare cancers, typically corresponding to an incidence of less than 15 per 100,000 people per year^30^. The rare tumors have a lower TLS than the non-rare tumors, and a comparison of the median of the two types of cohorts also reveals a significant difference (Supplementary Figure S2). Our hypothesis of lower TLS leading to better tumor recognizability can potentially account in part for this difference in incidence rates.

### Stratification of tumors by TLS is indicative of differences in immune infiltration and survival

Since the immune response will depend on the tumor’s immune environment in addition to T-cell recognizability, we investigated if there was any discernible pattern between the ‘selfness’ of a sample and the immune infiltration. There are established methods to perform immune cell deconvolution from bulk transcriptomic data, which we applied to the 10810 different samples across 32 different solid tumor cohorts in TCGA. Our workflow for this analysis (Figure 2A) involved (a) computing the TLS score for tumors in each cohort, (b) quantifying the difference in immune infiltration for each cell category for all possible stratifications into high and low thymus-like group. Our central question here was whether the ordering of tumors by TLS can be indicative of differences in immune infiltration.

**Figure 2.**
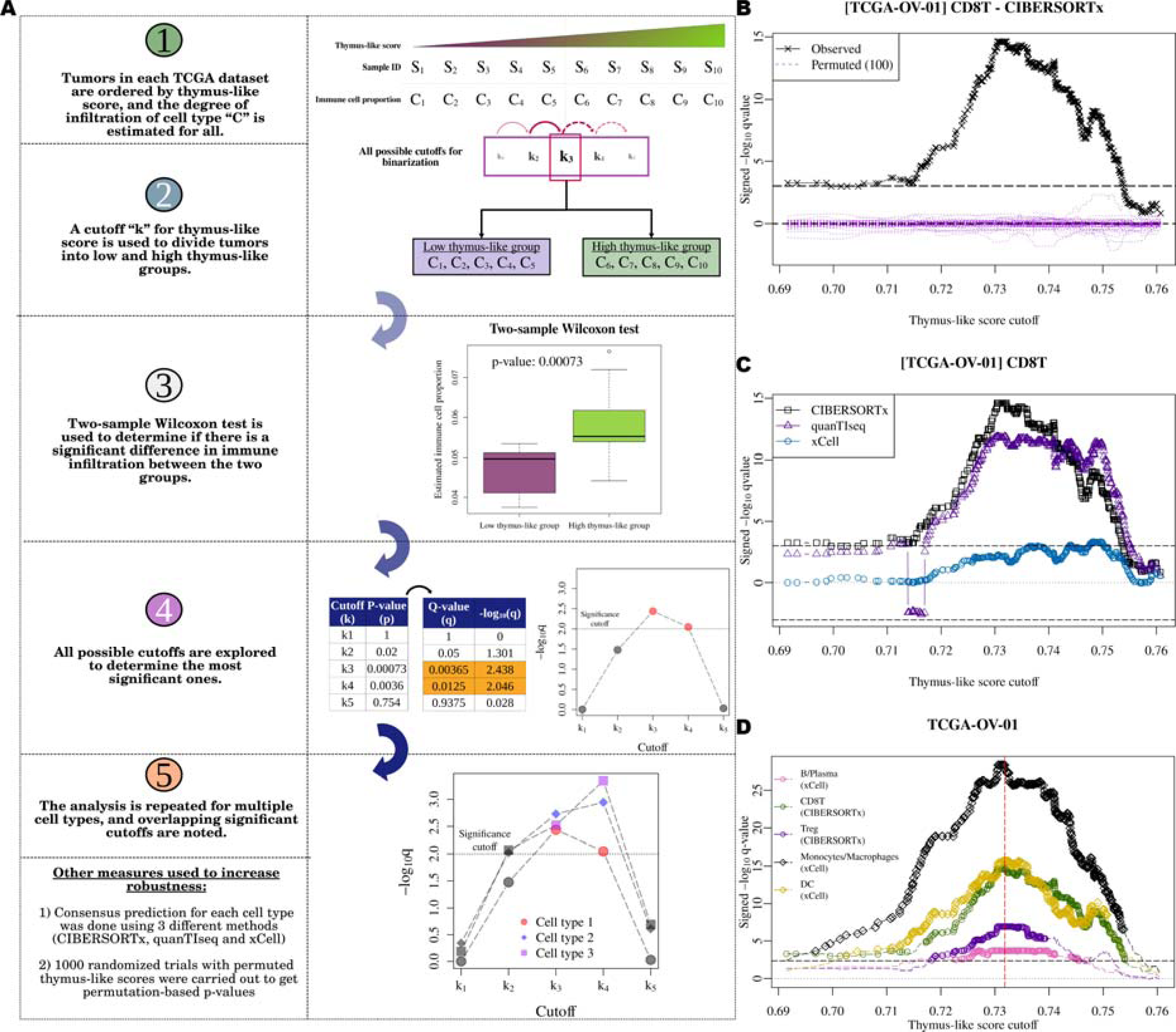
Schematic detailing the exhaustive stratification process along with representative plots. **(A)** Representative schematic showing the exploration of all possible stratifications. **(B)** Representative plot from the TCGA-OV cohort plotting the effect of choice of stratification cutoff on differential abundance of CD8 T-cells as estimated by a deconvolution method. The purple lines indicate the randomized control experiments. Only 100 are plotted here for the sake of clarity. The number of cutoffs that are significant for each cell category and deconvolution method across the different cohorts is listed in Supplementary Table 2 **(C)** Consensus plot for selecting significant stratification cutoffs for CD8 T-cells across three different deconvolution methods. **(D)** Final representative plot for all cell categories in a single cohort. Statistical significance between group-pairs was computed using the two-sample Wilcoxon test. The plots for all cohorts where at least one cell category is significant is shown in Supplementary Figure S3.

We first assessed immune infiltration of eight different cell categories by estimating their counts independently using three different immune deconvolution methods: CIBERSORTx, quanTIseq, and xCell^31–33^. Next, to identify the cutoff that leads to high-TLS and low-TLS groups having the maximum possible difference in terms of immune infiltration, we carried out a systematic and unbiased screening of all possible TLS cutoffs, since we had no prior information on the optimal stratification threshold in the context of immune infiltration, or if any such cutoff even exists. In brief, for every possible stratification of tumors into low and high TLS, we test whether there is a significant difference between the groups in terms of immune infiltration for specific immune cell categories predicted with each method using the Wilcoxon rank sum test. A representative plot for this stage of the analysis is shown in Figure 2B, where we plot the negative log of the q-value obtained for each cutoff. We add a sign to this value to indicate which of the two groups has higher infiltration, with positive values associated with increased cell abundance in the high thymus-like group. To test how significant the classification is, we performed 1000 randomization trials wherein we permuted the thymus-like scores and repeated the cutoff scanning procedure. The significance values obtained from this experiment act as a baseline for what is expected by random chance and allow us to calculate a second significance cutoff by recording the number of times the random trials resulted in more significant Wilcoxon rank sum tests than the observed ones. Next, we selected those cell categories that showed significant differences in perturbation between the stratified groups by at least two deconvolution methods (Figure 2C). For these cell categories, we heuristically find an optimal stratification cutoff for immune infiltration (Figure 2D). This is the point in the TLS scale that represents the statistically significant transition from low to high immune infiltration. To find a robust threshold, we used additional criteria that the low and high groups must each have at least ten samples each to reduce instability caused by small sample numbers. To account for multiple hypothesis testing, we use the False Discovery Rates (FDR) calculated from the p-values for testing significance.

We repeated this process for each TCGA solid tumor cohort and found that twenty-eight of the thirty-two cohorts could be stratified such that at least one cell category is differently abundant between them (Figure 3A and Supplementary Figure S3). The specific stratification cutoffs used for each cohort is reported in Supplementary Table 3. In most of these, the high thymus-like group had higher immune infiltration except for one dataset each for B/Plasma, CD4 T-cells, and NK cells. Monocytes/ Macrophages were the most commonly perturbed cell types, seen to be more abundant in the high thymus-like group in twenty-three datasets (Figure 3B). Similarly, the CD8 T-cells, regulatory T-cells, and Dendritic cell categories increased in eighteen, seventeen, and sixteen datasets, respectively. Overall, this points to a reliable increase in immune infiltration with an increase in the thymus-like scores within each TCGA cohort. While this appeared paradoxical at first, the results are clear and consistent in 9596 samples belonging to 28 tumor cohorts and point to a compensatory mechanism that attracts more immune cell infiltration into the tumor when the CD8 recognizability is low. Put together, this analysis clearly demonstrates that the TLS-based stratification divides tumors based on the extent of their tumor immune infiltration, a connection that is a new finding from this work.

**Figure 3:**
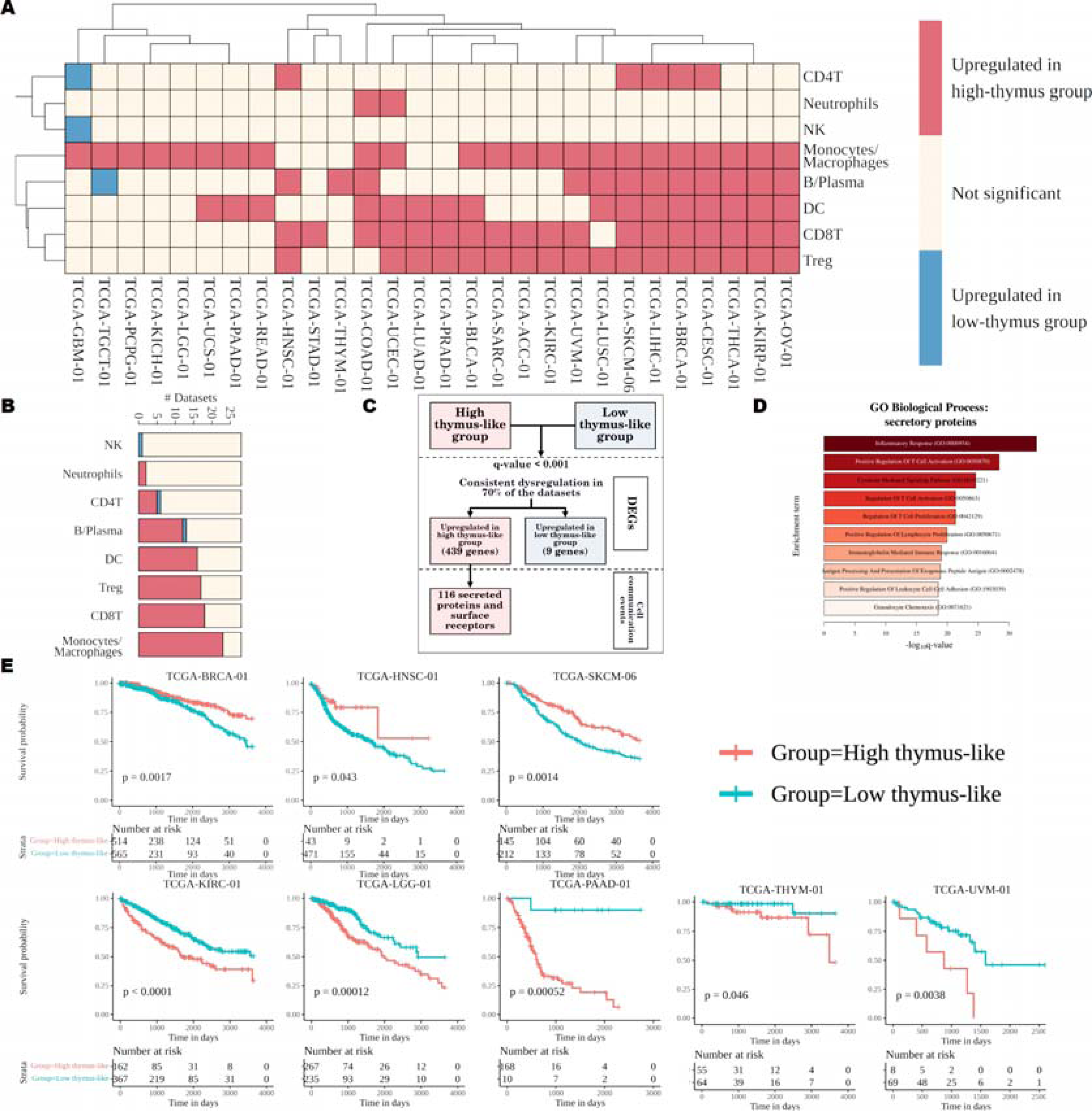
Stratification of TCGA cohorts by thymus-like scores reveals two kinds of tumor microenvironments. **(A)** Heatmap representing cell categories that are differentially abundant between the high and low thymus-like groups in each TCGA cohort. The color of the cell indicates the specific group in which a cell category is upregulated. **(B)** Barplot showcasing the number of cohorts in which each cell category is differently abundant. **(C)** Schematic explaining the meta-analysis of secretory proteins and receptors. **(D)** Top 10 GO Biological process enrichments of the meta-analysis. **(E)** Kaplan-Meier plots showcasing the different cohorts where the stratification process results in differences in survival.

### Gene expression meta-analysis reveals an immune attractive microenvironment in the high thymus-like groups

To ascertain whether the immune infiltration metrics are consistent with the ground reality at the molecular level, we conducted a meta-analysis to identify secretory proteins and receptors consistently dysregulated between the stratified groups. Specifically, we carried out a large-scale cohort-wise differential expression analysis of high and low thymus-like groups (Supplementary Table 4) and then identified genes that showed a consistent pattern of upregulation/downregulation across at least 70% of the cohorts through a random effects meta-analysis model. From these, we focused on only those genes whose proteins are either secretory proteins or surface receptors, using the list published as part of CellPhoneDB as reference^34^. These proteins provide a small window into the cell-cell communication events that are differential in the tumor microenvironments of each group. The meta-analysis identified 116 secreted proteins and receptors consistently upregulated in the high thymus-like group across all TCGA tumor cohorts (Figure 3C, Supplementary Table 5) (1.5 fold dysregulation, q-value < 0.01 and sign consistency in 70% of the cohorts). Enrichment analysis of these genes revealed upregulation of general inflammatory response, chemotaxis related genes, lymphocyte proliferation and activation, and genes of the antigen presentation pathway (Figure 3D). Specific analysis of the chemokines revealed chemo-attractiveness to various cell types, with T lymphocytes being the most targeted (Supplementary Table 6). The upregulated interleukins have a pleomorphic effect and are all known to affect T lymphocytes. HLA genes of both class I and II were also seen to be upregulated, along with co-stimulatory molecules involved in providing the second signal for activation of mature, but naive T lymphocytes. In addition to these, secretory proteins that upregulated these co-stimulatory factors on the antigen presenting cells were also seen to be upregulated. Various T-cell checkpoints pairs were also upregulated. Taken together, the enrichment results suggest that the high thymus-like group is enriched for activation of naive T cells, and concomitantly, is associated with the machinery that keeps unregulated T cell activation in check. While the low thymus-like group is expected to be better recognized by T-cells, the tumor microenvironment is more immunologically active in the high thymus-like tumors.

### The two facets of tumor immune response: Recognizability and Immune infiltration

Both immune infiltration and tumor recognizability are important determinants of successful T-cell response to tumors, and each of the stratified groups is associated with only one of these two facets. We therefore wanted to assess if one of these facets provided a larger advantage than the other. Our analysis indicated that 8 tumor cohorts had significant survival differences between the stratified groups (Figure 3E). The low-thymus group showed better survival in 5 cohorts, these being the Kidney renal clear cell carcinoma (KIRC), Low grade glioma (LGG), Glioblastoma (GBM), Pancreatic adenocarcinoma (PAAD), thymoma (THYM) and Uveal Melanoma (UVM) cohorts. Interestingly, 3 tumor cohorts (Breast invasive carcinoma - BRCA, Head and Neck squamous cell carcinoma - HNSC and metastatic Skin Cutaneous Melanoma - SKCM) had an increased abundance of CD4 T-cells, and showed better survival in their high thymus-like groups indicating that the poorer recognizability may in some way provide a feedback to inform the extent of infiltration so as to result in higher CD4 immune-infiltration as compensation for the lack of recognizability due to high thymus-likeness. The tissue-of-origin of these tumor cohorts are also associated with higher thymus-like scores (Figure 1B) than the former 5 cohorts, possibly indicating different requirements for successful tumor clearance depending on the baseline level of surveillance of each tissue. Of the 5 cohorts where lower thymus-like scores lead to better survival, LGG, GBM and UVM originate from tissues that have less immune surveillance.

## Discussion

For a successful immune response against tumors, immune cells are required to recognize the tumors and mount an immune response of sufficient magnitude against them. This relies on the ability of immune cells to recognize the tumor cells as aberrant and mount a robust response to eliminate them. Tumors on their part, can use either or both of these aspects to evade destruction by the immune system while the immune cells continuously seek out and eliminate cancerous threats, akin to a game of hide-and-seek. The primary strategy for the immune cells to achieve their goal is to recognize a tumor cell as aberrant with high specificity such that their targets are restricted to tumor cells without harming the normal cells. The key components of the process then are recognizing an antigen representing an aberrant structure on the tumor cells as well as distinguishing self from non-self. Two types of aberrations can be found in tumor cells - (a) an abnormal protein or molecular complex serving as a neo-antigen (TSAs) and (b) an abnormal level of a normal protein that by some as yet unknown mechanism attracts the immune cells (TAAs). In both cases, the basic paradigm that the cells employ is to distinguish self from non-self. While it is easier to comprehend how the neo-antigens are perceived as non-self and hence get recognized especially by the HLA molecules, there is much less clarity on what precisely comprises the self vs non-self axis in the case of TAAs. To the best of our knowledge, there has been no systematic treatment of this problem.

Our study addresses this gap and shows that the comparison with the thymus presents an opportunity to define selfness and non-selfness more precisely. It is possible that aberrantly expressed normal proteins in tumor cells can be processed and presented as self-antigens by the HLA molecules. While self-antigens are typically tolerated by the immune system, aberrant expression or presentation of self-antigens in the context of tumor cells may trigger immune recognition and activation through mechanisms of surveillance of cellular homeostasis and detection of abnormal cellular stress.

In our study, we model the recognizability of tumors to T-cells using the thymus-like score and surprisingly find that recognizability and immune infiltration are inversely correlated. This correlation could be due to the elimination of highly infiltrated tumors easily recognizable to T-cells at an earlier stage of tumorigenesis. This could mean those easily recognizable tumors must either inculcate an immune-cold microenvironment, or they would have to alter their transcriptional program to become less recognizable to survive the immune onslaught. Specifically, our analysis reveals an increase in the abundance of CD8 T-cells, T-reg cells, dendritic cells, monocytes and macrophages with a rise in the TLS in at least half the tumor cohorts analyzed. In terms of survival, tumors that arise from tissues with lower TLS are associated with better survival if the tumors have low thymus-like scores themselves, but the opposite is seen in cases where tumors originate from tissues with high TLS and where the group with high TLS also has increased CD4 T-cell infiltration. Inherent differences in immune infiltration and TLS of the tissue of origin for the tumors could be indicative of the type of immune response required to clear the tumors.

Our analysis draws the need for future experiments to closely examine the link between the recognizability of tumors and the observed clinical outcome. Some examples include analyzing the response of identical mice to tumors that differ by their thymus-like score or, alternatively, manipulating the expression profile of mTEC^hi^ cells in different mice and observing the immune response to identical tumors.

In addition, the results we present can easily be seen to have implications for utilizing different therapies based on where the recognizability/infiltration spectrum of the tumor falls. Tumors that are highly recognizable but poorly infiltrated would benefit from agents that enhance immune infiltration. In contrast, those not very recognizable but well-infiltrated would be best complemented by drugs that alter the tumor profile to be more recognizable. Drugs that push the transcriptional profile of tumors to be dissimilar to that of mTEChi cells could be a valuable addition to chemotherapy regimens to enhance the activity of T-cells.

## Materials and Methods

### Identification and processing of mTEC^hi^ datasets

NCBI’s Gene Expression Omnibus database was screened for datasets from studies profiling the mTEC^hi^ cell subset isolated from pediatric surgery patients (PRISMA chart is shown in Supplementary Figure S1A)^21,22,35^. GSE176445 and GSE201719 mTEC^hi^ samples were retrieved from NCBI’s SRA database using SRAToolkit 3.0.0^36^. TrimGalore 0.6.7 was used to remove adapters. STAR 2.7.10a and RSEM 1.3.3 was used to quantify gene expression using the ENSEMBL Human genome (release 106) as the reference^37–39^. Genes were ranked sample-wise according to their TPM values, with the highest TPMs get the best ranks (smallest rank values). We obtained the final representative rank values (mTEC^hi^) for each gene by ordering the mean ranks of each gene across all samples. The mean reference vector correlates well with all samples (Supplementary Figure S1B).

### Calculation of the ‘thymus-like score’

We define the thymus-like score for any sample as the Spearman correlation between that sample’s ranks of TPM values against the mTEC^hi^ rank vector described previously. In case of ties between multiple genes, we assign them the maximum possible rank. This ranking system ensures that 0 TPM is associated with the same rank across all samples. We determined beforehand the set of genes whose expression data was available across all samples in the analysis and used only those for the thymus-like score calculation. The general schematic is shown in Figure 1A.

### Processing of Human Protein Atlas data

The consensus transcriptome dataset with 54 tissues was downloaded from the Human Protein Atlas^26^, and their TLS were calculated. The analysis uses the normalized TPM values reported for these tissues. The sorted TLS were fed to Cksegs.1d.dp function from the Ckmeans.1d.dp (4.3.4) package to carry out univariate segmentation analysis^40^. In some cases like the brain, the dataset contains 12 different tissues sourced from the brain: Amygdala, Basal ganglia, Cerebellum, Cerebral cortex, Hippocampal formation, Hypothalamus, Medulla oblongata, Midbrain, Pons, White matter, Thalamus, and Choroid plexus and we calculate the mean thymus-like score of these tissues. We show that this simplification is justified, as these tissues were found to share highly correlated transcriptomic profiles (Supplementary Figure S1D).

### Processing of TCGA data

Transcriptomic and clinical data from the TCGA project was downloaded using the GenomicDataCommons 1.22.3 R package. We filtered for all transcriptomic STAR counts data from patients with clinical data available, with the additional filter that the samples are either solid tumors (primary, recurrent, or metastatic) or non-tumors from patients with solid tumors. Sample redundancy was then removed by selecting only one vial code for each previously mentioned sample type per TCGA dataset. The filtering resulted in the selection of 10810 transcriptomic samples, including 9670 primary tumors, 43 recurrent tumors, 392 metastatic tumors, and 705 non-tumor samples. (Supplementary Table 1 and Supplementary Figure S1E).

### Immune deconvolution

CIBERSORTx, quanTIseq, and xCell were used for estimating the abundance of immune cell types^31–33^. quanTIseq and xCell were run through the immunedeconv (2.1.0) wrapper R package, which in turn calls the quantiseqr (1.6.0) and xCell (1.1.0) packages^41^. CIBERSORTx absolute mode was run using the “no.sumto1” method using the LM22 signature matrix via the Docker program provided by the authors after registering for a user token. The cell types produced by different methods were unified into eight categories for simplifying the downstream analyses: B/Plasma cells, Myeloid Dendritic Cells, CD8 T cells, Non-regulatory CD4 T cells, Regulatory T cells, Monocytes/Macrophages, Natural Killer cells, and Neutrophils. In cases where a method provides a more detailed split of one of these categories, they were summed up to provide the unified category mentioned above.

### Consensus cutoff selection for one cell category

The consensus cutoff scanning was done individually for each tumor cohort. In cases where the tumor cohort had solid tumors of multiple types (primary, recurrent or metastatic), the analysis was conducted separately for each type. Only cohort subdivisions that contained at least 20 tumor samples were considered for the analysis, resulting in the analysis of primary tumors from all cohorts and the metastatic cohort from the SKCM cohort.

Any specific stratification of tumors into high and low thymus-like groups was considered to show significant differences in immune cell counts if the q-values generated using Wilcoxon rank sum tests was less than 0.005 and if less than 5 of the 1000 randomization trials produced equal or better significance values. Consensus was obtained across deconvolution methods by requiring that each stratification cutoff pass this significance filter when two different deconvolution methods were used, with both methods agreeing on the direction of perturbation between the stratified groups.

To prevent a single deconvolution method from dominating the average significance values, we perform an optimal univariate K-means clustering of the negative log_10_ q-values from the Wilcoxon rank sum tests for each method and then rank each cluster based on their significance, resulting in cutoffs providing similar significance values being grouped together. This value is then averaged across the different deconvolution methods. This results in lower values for stratification cutoffs that provide the best separation across all deconvolution methods. Again, for increased robustness, the cluster rank average is taken for the worst two significant ranks across the methods.

### Consensus cutoff selection across all cell categories

We use a heuristic to guide the selection of a single cutoff across cell categories within a single cohort that maximizes the number of cell categories that are perturbed between the high and low thymus-like groups, while maintaining as high a significance value as possible in the Wilcoxon rank sum tests across the different deconvolution methods.

First, we catalog the number of cell categories that can be stratified to show significant differences between the high and low thymus-like groups. The stratification points for these categories may or may not overlap. We select only cutoffs such that at least 60% of the significant cell categories are stratified into differentially abundant groups. This is relaxed only in cases where there are only 2 cell categories that are significant. Then, these cell categories are ranked based on the mean of cluster rank average for each significant cell category. Finally, to break ties, we use the mean of the actual negative log_10_ q-values from the Wilcoxon rank sum tests. For the case of TCGA-LGG and TCGA-PRAD however, we chose only the latter category to eliminate cases where multiple borderline significant cell categories influenced the results, thereby resulting in a more robust cutoff selection.

### Differential expression analysis and gene enrichment

TCGA STAR counts data was imported into R using tximport 1.26.1 and differential expression analysis was carried out cohort-wise between the high and low thymus-like groups using limma 3.54.2^42,43^. The results from the cohort-wise analysis was taken forward for meta-analysis using the MetaVolcanoR package 1.14.0. Genes with random summary statistics of 1.5 fold change with q-value <= 0.001 and showing sign consistency in at least 70% of the datasets were considered to be consistent DEGs between the two groups. This list was then filtered for secreted proteins and membrane receptors using CellPhoneDB as a reference^34^. Finally, GO Biological Process Enrichment was carried out using EnrichR^44^.

### Survival analysis

Log-rank test was used to compute significance p-value for survival differences between the high and low thymus-like groups. Cohorts which showed significant survival differences were visualized using Kaplan-Meier plots. The R packages survival 3.5 and survminer 0.4.9 packages were used for these^45,46^.

## Supporting information

All supplementary tables

## Acknowledgements

We acknowledge support from the Bioinformatics grant, Department of Biotechnology, Government of India (BT/PR40186/BTIS/137/3/2021). The results shown here are in whole or part based upon data generated by the TCGA Research Network: https://www.cancer.gov/tcga. Naren Chandran Sakthivel is funded by the Council of Scientific & Industrial Research, Government of India (Senior Research Fellow - File no: SPM-07/079(0287)/2019-EMR-I).

## Code Availability

The code used for this analysis can be accessed and downloaded from: https://gitlab.com/narencs179/thymus-like-score

## Data Availability

Processed data for this analysis can be accessed and downloaded from: https://doi.org/10.5281/zenodo.10829212

## Author Contributions

The initials used for the author contributions are as follows:

- Naren Chandran Sakthivel – N.S.
- Anoushka Chinmayi – A.C.
- Nagasuma Chandra – N.C.

**Conceptualization –** N.C. and N.S.; **Methodology –** N.C. and N.S.; **Software and Investigation –** N.S. and A.C.; **Data Curation –** N.S. and A.C.; **Writing – Original Draft –** N.C. and N.S.; **Writing – Review & Editing –** N.C. and N.S.; **Supervision –** N.C.

## Supplementary Information

**Supplementary Figure S1:**
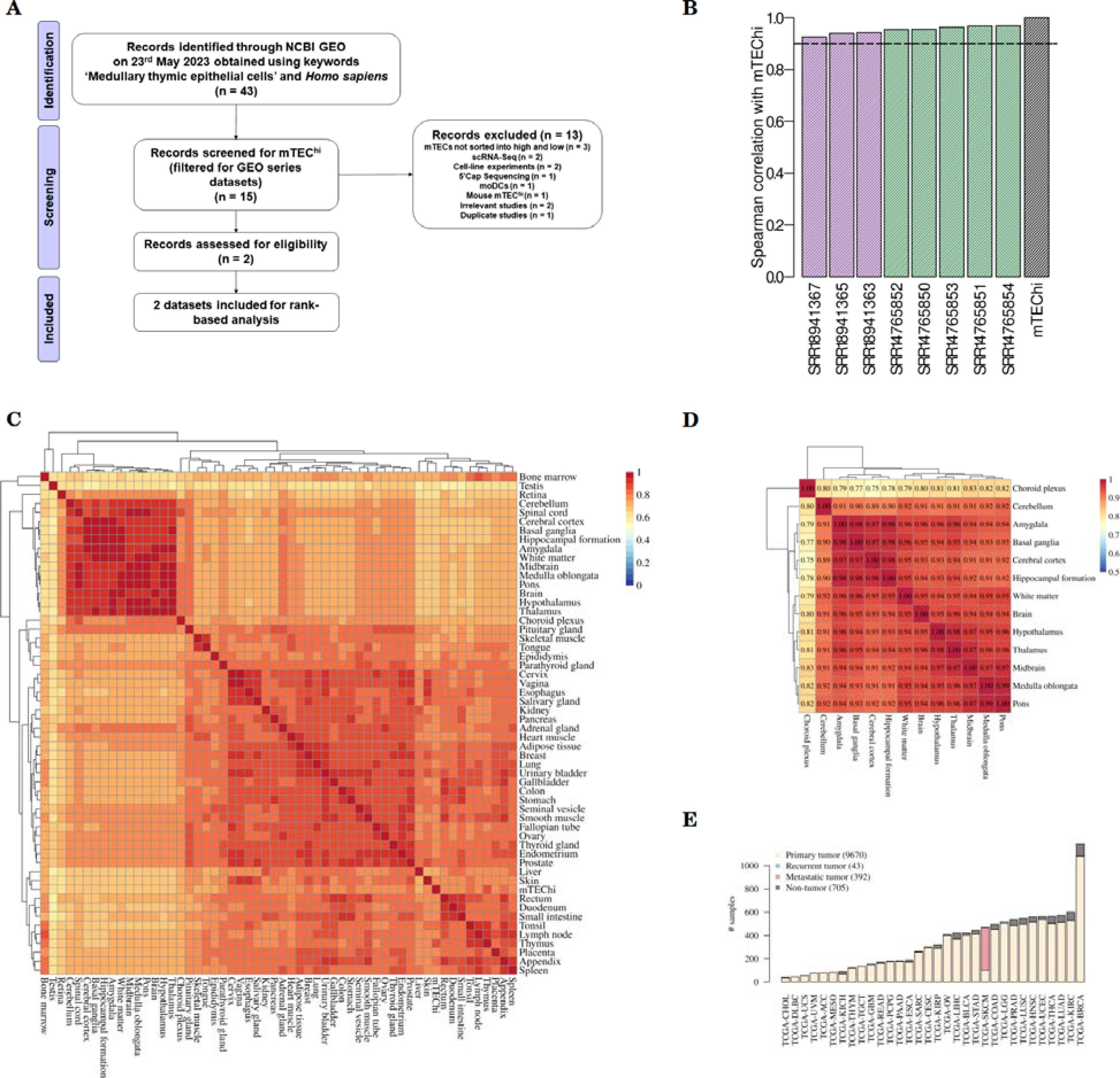
**(A)** PRISMA flow diagram for mTEC^hi^ dataset search. **(B)** Spearman correlation of different mTEChi samples from SRA against the mTEChi mean rank vector calculated from them. The color of each bar denotes its dataset of origin. While there is a dataset-specific difference, the correlation is greater than 0.9 (dotted line) for all samples. **(C)** Spearman correlation of all transcriptomic profiles from Human Protein Atlas and the mTEC^hi^ reference profile used in the study. Two broad clusters are seen, with the smaller cluster being enriched for immune privileged tissues: testis, retina and different regions of the brain. The rest of the tissues form one large cluster, with the mTEChi reference profile clustering within this group. **(D)** Spearman correlation of transcriptomic profiles from different brain tissues in the Human Protein Atlas indicates high correlation between them. The representative computed from these regions (labelled as ‘Brain’) has a high correlation (≥ 0.91) with all regions barring the choroid plexus, which still has a high correlation(0.8). **(E)** Distribution of samples across different TCGA datasets: The number of samples ranges from 44 in the Cholangiocarcinoma (TCGA-CHOL) cohort to 1186 samples in the Breast Invasive Carcinoma (TCGA-BRCA) cohort. The majority of the samples in each cohort is composed of primary tumor samples, with the exception of the Skin Cutaneous Melanoma (TCGA-SKCM) cohort, where metastatic samples predominate.

**Supplementary Figure S2.**
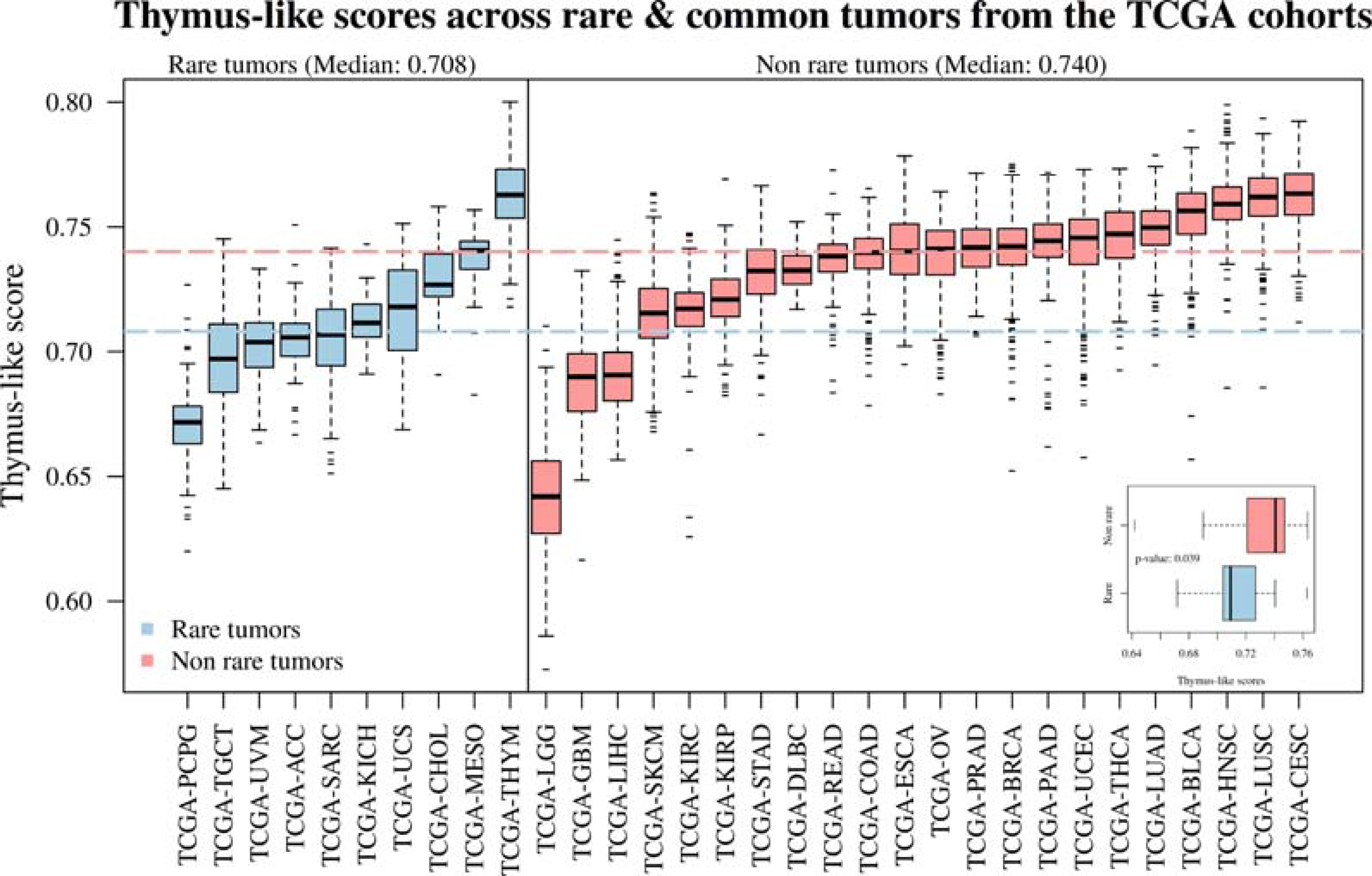
Distribution of TLS across rare and non-rare tumors. The thymus-like scores of the different TCGA cohorts are arranged according to their rarity. The median scores of the rare and non-rare tumors are shown by the blue and red dashed lines, respectively. The subplot in the bottom right compares the median thymus-like scores of the rare cohorts with the non-rare cohorts. Significance value was calculated using Wilcoxon rank sum test.

**Supplementary Figure S3.**
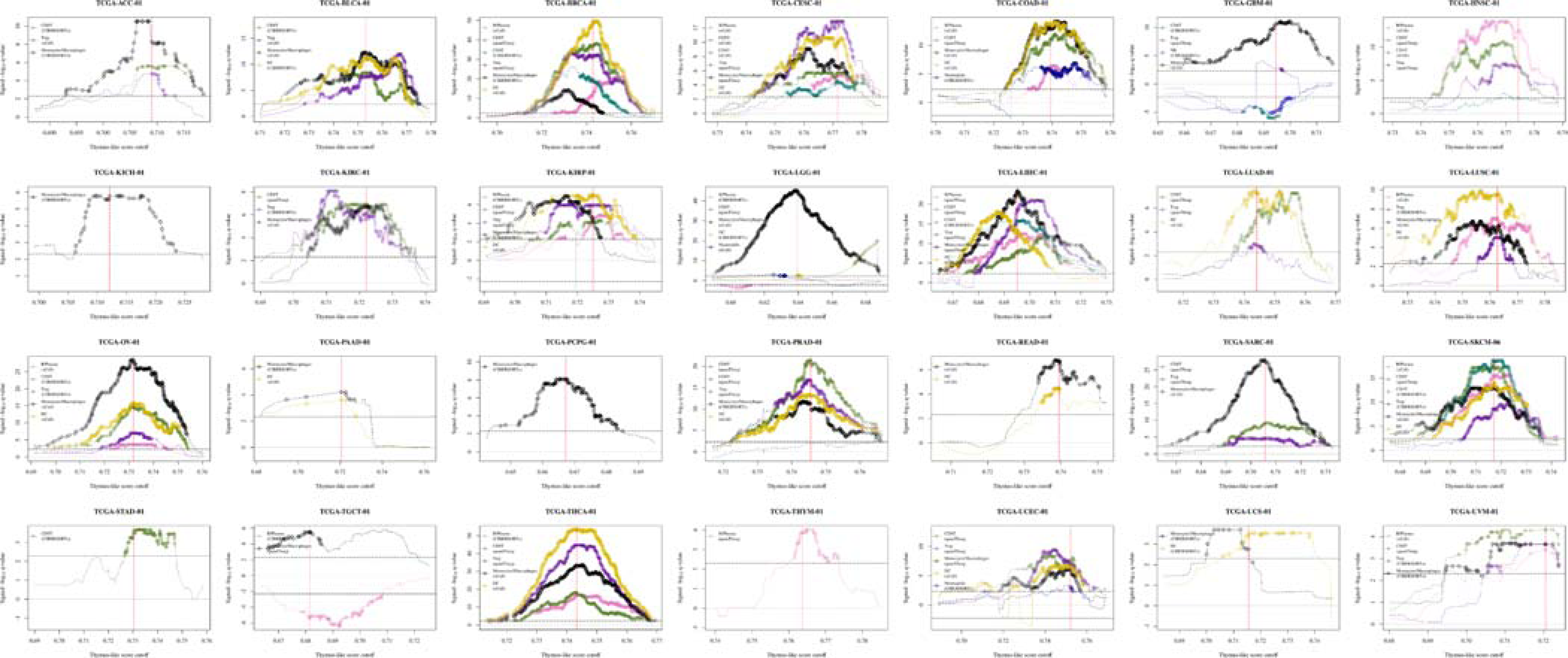
Optimal stratification cutoff selection across 28 cohorts. For 28 TCGA cohorts, there exists an optimal stratification cutoff wherein tumor samples split into high and low TLS groups show significant difference in at least one cell category. The plots here show the specific cutoffs selected for each of these cohorts. Each cell category is represented by one method here. However, every significant cutoff showcased here is significant by at least one other method also. The red line marks the optimal stratification cutoff selected by the heuristic method.

